# Within-subject daily rhythmicity and longitudinal stability of hunger, satiety, and physiological markers

**DOI:** 10.1101/2025.10.10.681393

**Authors:** Xi Wang, Abhishek S. Prayag, Ni Tang, Yanlong Hou, Marc Thevenet, Claude Gronfier, Tao Jiang

## Abstract

Hunger and satiety are interoceptions involved in the control of food intake. How they change across the day and from one day to the next, as well as their relationships with physiological parameters, remain poorly understood.

In a five-day laboratory study, twenty healthy male participants (24.2 ± 3.3 years) were given three meals per day (breakfast, lunch, dinner), at times based on each individual’s internal time. Subjects had an 8-h sleep opportunity at night recorded by polysomnography, during which time they were exposed to 4 different light intensities (0, 3, 8, or 20 lux). Hunger and satiety were assessed before and after each meal. Heart rate was measured continuously and glucose levels were sampled every 15 minutes.

We found that: 1) Hunger and satiety varied dynamically with the lowest levels at breakfast and the highest at lunch and dinner (*p* < 0.01); 2) the intensity of hunger and satiety remained stable across the five experimental days, and were subject-dependent; 3) Heart rate was higher after than before meals (*p* < 0.0001), and higher around breakfast than dinner; 4) Glycemia before and after meals was respectively stable throughout the day and over the study period; 5) Interoception was correlated with fluctuations in heart rate, glucose levels and sleep duration. Overall, our results reveal that hunger and satiety perception are individual characteristics.

To better understand the role of interoception in food intake control, both hunger and satiety should be measured, but separately in their respective states.

## 1. Introduction

Food intake is a critical behavior for living organisms because it satisfies energy needs and maintains internal homeostasis. Interoception, the process by which internal physiological states (e.g., the body’s energy status) are sensed and interpreted, plays a central role in regulating appetite status and controlling the behaviors necessary for establishing and maintaining internal balance. The two main interoceptions related to food intake are hunger and satiety. They have long been considered as the most important biological drivers of eating behavior. Hunger is the body’s natural response to energy deficiency, and is characterized by sensations such as a pit in the stomach, gurgling, or cramping. These sensation signals to the brain that the body is running low on energy and requires calories. In contrast, satiety reflects a state of fullness, that develops gradually during the eating course and peaks 20-30 minutes afterward (Blundell et al., 2010; Green et al., 1997; Holt et al., 1995; Mela, 2006). As satiety progresses, the motivation to eat decreases until food intake ceases, reflecting energy repletion (Bellisle, 2005; Woods et al., 1998). The alternation between hunger and satiety is influenced by both physiological processes as well as external factors, such as food products, cognition, socio-culture, and the environment, which determine the choice, amount, timing, frequency and rhythm of food intake (Bellisle, 2005; Berthoud et al., 2020; Broussard et al., 2016; Kabir et al., 2018; Marcelino et al., 2001; McNeil et al., 2017; Paoli et al., 2019). However, the temporal variation of hunger and satiety over the course of a day, as well as over several consecutive days, remains underexplored. Furthermore, the interactions between these sensations and physiological markers, such as heart rate and glucose levels, are unclear.

The fluctuation of hunger and satiety around a meal is well documented. Hunger peaks before a meal and then decreases and disappears afterward, while satiety does the opposite. Several chronobiological studies have examined these fluctuations over the course of a day and have demonstrated that they are rhythmic. Depending on the study design, hunger peaks in the evening (Sargent et al., 2016) or in the morning (Qian et al., 2019; Scheer et al., 2013), and satiety peaks at night (Sargent et al., 2016). While the rhythmicity of hunger and satiety has been documented, few studies have examined these sensations multiple times (meals) per day and over several days in the same individuals, and under an ecological protocol. This knowledge gap is particularly relevant in an obesogenic environment where constant food availability increasingly emphasizes exteroception (external sensory cues) over interoception. As exteroception plays a more dominant role in controlling food intake in an obesogenic environment, the precise contribution of interoception to controlling food intake becomes even less clear, and thus warrants further investigation.

The perception of hunger and satiety may be different between individuals, since some people base their perceptions on overall discomfort, while others base them on stomach fullness. Objective physiological measures may be thus useful for understanding the nature of these perceptions. Although heart rate (HR) reflects autonomic nervous system activity (Jose & Collison, 1970), the relationship between HR and hunger and satiety measures has rarely been explored.

Glucose levels are an indicator of energy availability and are closely related to hunger and satiety. Decreases in blood glucose levels before a meal are often associated with the onset of hunger, and increases in glucose levels after a meal contribute to the onset of satiety (Campfield et al., 1985; Campfield & Smith, 1990a; Louis-Sylvestre, 1976). Understanding the interplay between glucose dynamics and hunger or satiety sensations for different meals (e.g., breakfast, lunch and dinner) could provide valuable insights into the interaction between metabolic signals and interoception. For instance, some studies suggest that glucose responses are more pronounced after breakfast than after other meals, possibly due to circadian influences on glucose metabolism (Van Cauter et al., 1997). However, the relationship between glucose patterns and hunger or satiety at different times of the day remains unclear, especially with repeated daily assessments.

Hunger and satiety are also both closely linked to sleep. Disruptions in sleep quality or duration can modify the perception of hunger and satiety. This can lead to increased resting energy expenditure (Markwald et al., 2013), an altered food preference for energy-dense foods (McNeil et al., 2017), and greater overall food consumption (Brondel et al., 2010).

Additionally, light as the primary cue that synchronizes the circadian clock and the sleep-wake cycle, affects many physiological functions (Prayag et al., 2019). Nevertheless, the potential influence of nocturnal light exposure on food intake through interoception (hunger and satiety) remains unexplored. The current study aimed to investigate: 1) characteristics of interoception (hunger and satiety) related to food intake and how they evolve over one day and several days, and 2) physiological patterns, such as heart rate and glucose levels, in relation to the hunger and satiety sensations. 3) the effect of low-intensity light exposure at night on interoception the next day and whether this effect is secondary to light-induced sleep duration.

## 2. Material and Methods

### 2.1. Participants

As the menstrual cycle has been shown to impact appetite and energy intake (Barbosa et al., 2015; Candan et al., 2025; Gorczyca et al., 2016; Krishnan et al., 2016; Rogan & Black, 2022; Tucker et al., 2025; Yukie et al., 2020), metabolic rate (Benton et al., 2020; Solomon et al., 1982), sensory perception (Stanić et al., 2021; Watanabe et al., 2002), sleep, and circadian rhythms (Jeon & Baek, 2023), only male participants were included in this study to minimize its potential confounding effects.

Twenty healthy male participants (age: 24.2 ± 3.3 years) were recruited for a five-day inpatient laboratory study. Each participant completed several questionnaires on personal information, general health, and sleep quality (Pittsburgh Sleep Quality Questionnaire, PSQI, Buysse et al., 1989); chronotype typology (Horne & Ostberg, 1976); the Munich Chronotype Questionnaire (MCTQ, Roenneberg et al., 2003); the Beck Depression Inventory (BDI, Beck et al., 1961); and the Eating Attitude Test (Garner & Garfinkel, 1979).

The inclusion criteria for participants were: PSQI score ≤5, Horne & Ostberg score between 31 and 73, BDI score ≤10, and Eating Attitude Test score ≤20. Participants were excluded if they had performed shift work or traveled across time zones in the previous two months. They were instructed to maintain a consistent sleep/wake schedule for an average of two weeks prior to the laboratory session, with a deviation of less than 30 minutes from the target bed and wake times. This was verified using both a daily sleep diary and an actimeter (ActTrust, Condor Instruments, São Paulo, Brazil) worn on the non-dominant wrist. Participants also completed light and food exposure diaries.

### 2.2. Study design

Participants were admitted to the laboratory on Monday and were discharged on Friday evening (see Figure 1A).

**Figure 1.**
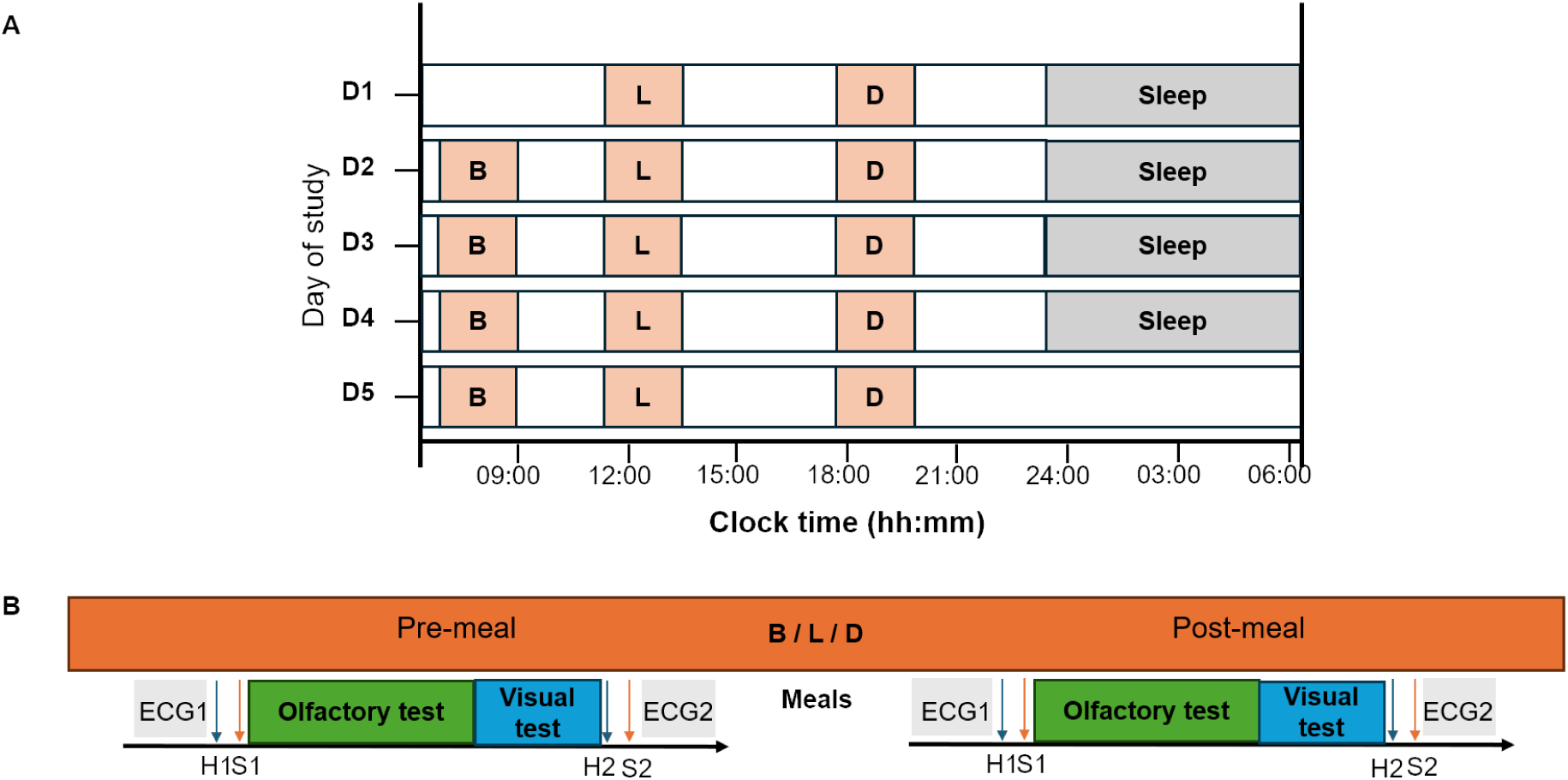
Illustration of the 5-day experimental protocol in the laboratory. **(A)** Subjects entered the laboratory on Monday, two hours after their habitual waketime, and were discharged on Friday at their habitual bedtime. Three meals were provided each day (except Monday because subjects arrived in the laboratory after their breakfast at home): breakfast (B), lunch (L), and dinner (D). Sleep times were scheduled at each subject’s habitual sleep times (here, bedtime and wake time are set at 2330 and 0730, respectively). **(B)** The pre-meal sensory tests were conducted approximately seven minutes before breakfast and lasted eight minutes. Before lunch and dinner, the tests started approximately 24 and 50 minutes before the meals, respectively, and lasted about 13 minutes each. Participants started with a 1-minute electrocardiogram (ECG1) measurement, followed by an assessment of hunger (H1) and satiety (S1). Next, participants underwent olfactory and visual tests. At the end of the test, a new self-evaluation of hunger (H2) and satiety (S2) was performed, and a final one-minute ECG-2 was taken. The post-meal sensory tests were performed 20 minutes after the end of meal and followed the same procedure as the pre-meal measurements.

#### Experimental and light conditions

During the study, participants were in a time-isolation conditions (deprived of all-time cues, such as natural light, watches, clocks, TV, computers, smartphones, and radios), and were not allowed to lie down in bed or take a nap during the day. Physical activity was limited to walking around the room and stretching, in order to avoid significant increases in body temperature (more than 0.5°C).

Light intensity was controlled both during the day and at night. During the day, from awakening until bedtime, the light intensity was maintained at 90 lux (average angle of gaze). At night, from bedtime to wake time (eight hours), low artificial light at night was set to one of four light conditions: 0, 3, 8, or 20 lux. Each participant was exposed to all four light conditions (one light condition per night). The order of light presentation was counterbalanced according to a 4×4 Latin square design.

#### Individual scheduling

Sleep, mealtimes and all other measures and recordings were calculated and scheduled individually, based on each participant’s habitual sleep-wake schedule at home. This allows to schedule events at times that correspond to the same internal time across subjects (Gronfier et al., 2007). Habitual sleep times were calculated for each subject using the following procedure: First, we averaged the bedtimes and wake times of the last seven sleep episodes documented by each subject in their sleep diary and verified with actimetry. Second, we found the mid-sleep time between the average bedtime and wake time. Third, we subtracted four hours from the mid-sleep time to obtain the average habitual bedtime, and added four hours to the mid-sleep time to obtain the average habitual wake time (Daguet et al., 2022). Mealtimes were set at the habitual wake time (HWT) plus 50 minutes for breakfast, plus 320 minutes for lunch, and plus 720 minutes for dinner (corresponding to 7:50 a.m., 12:20 p.m., and 7:00 p.m., respectively, for who habitually wakes up at 7:00 a.m.).

#### Meals

For breakfast (B), participants selected their first laboratory breakfast from a predefined list and were required to maintain this, choice for consecutive mornings. Available options s included dairy products (milk, yogurt, and cheese), fruit juices (apple and orange), fresh fruit (apples and oranges), bread, crispbread, butter, jam, cereal, and caffeine-free coffee.

Lunch (L) and dinner (D) were provided by the hospital restaurant, which had rotating menus on a quarterly basis with quasi equal calories. A typical meal included a starter of either a green salad with dressing or a mixed vegetable salad (tomato, potato etc., with dressing); a main course of meat, fish, or eggs with vegetables; a side dish (e.g., rice or noodles with bread); a dessert (e.g., an orange, a banana, an apple, a fruit salad, a chocolate mousse, a compote), and a dairy product (e.g., cheese, yogurt, or pudding).

The meals were tailored to the participants’ age group and nutritional needs. They were designed to provide an average daily caloric intake of ∼2430 kcal per day, distributed as follows: ∼55% carbohydrates, ∼15% protein, and 30% fat.

The estimated average caloric intake per meal was ∼600 kcal for breakfast, 915 kcal for lunch and 915 kcal at for dinner. Breakfast’s energy distribution was as follows: ∼90 kcal from fruit juices, ∼170 kcal from dairy products, ∼190 kcal from cereals, bread or crispbread, ∼35 kcal from fresh fruits, ∼85 kcal from butter and ∼30 kcal from other food items. Lunch and dinner followed a similar structure, with an additional dairy product served at dinner. The calorie contributions were approximately ∼40 kcal for a mixed salad and vinaigrette, ∼540 kcal for the main course, ∼200 kcal for the side dish, ∼65 kcal for the dairy product, and ∼70 kcal for dessert. Participants were informed that they could stop eating when they felt full. Each plate was weighed before and after each meal. Participants finished almost all of their food at each meal.

### 2.3. Measurements of hunger and satiety and sensory tests

Participants completed six experimental sessions each day, corresponding to the periods before and after breakfast (B), lunch (L), and dinner (D). This schedule was repeated throughout the five-day study (Figure 1A).

Each testing session began with a one-minute baseline electrocardiogram (ECG) recording. During this time, participants were asked to remain calm and motionless while their heart rate (HR) was recorded. Immediately afterward, participants completed self-assessments of hunger and satiety using a computerized VAS (Figure 1B).

Participants then performed both olfactory and visual sensory tests. The olfactory test consisted of rating the three attributes (liking, wanting, and disgust) on a visual analog scale (VAS) for each of the 14 food odors. The visual test consisted of rating the same three scales after viewing a picture corresponding to the food odors presented in the olfactory test. All sensory tests and instructions were provided via SuperLab software (Cedrus, San Pedro, CA, version 5.0). Throughout testing, participants sat 60 cm in front of the computer screen and communicated with the experimenter via interphone.

The pre-meal sensory sessions began approximately 15 minutes before breakfast (B) and lasted eight minutes. Before lunch (L) and dinner (D), the tests began 24 and 50 minutes before the meal, respectively, and lasted 13 minutes. The same sensory tests were performed an average of 28 minutes after each of the three daily meals (see Figure 1B).

### 2.4. Measures

#### 2.4.1. Hunger and satiety

Participants rated their hunger and satiety levels using a visual analog scale (VAS). A 30-cm-long line was displayed on the screen with the left end labeled with “not hungry at all”, or “not satiated at all”, and the right end labeled with “extremely hungry” or “extremely satiated.” Participants were asked to click on the line that corresponded to their level of hunger or satiety. The click points on the VAS were converted to numerical values ranging from 0 to 15, with 0 representing the leftmost point and 15 representing the rightmost point.

#### 2.4.2. Heart rate

Heart rate (HR) was recorded at a frequency of 256 Hz using a polysomnography (PSG) monitoring system (Vitaport-4 digital recorder, TEMEC Instruments, Kerkrade, Netherlands). Two electrodes were placed: one at the right infraclavicular fossa, and one at the V6 position. As shown in Figure 1B, HR was recorded from one minute before the first hunger assessment (H1) to one minute after the second satiety assessment (S2), for both before (pre) and after (post) each meal. The mean HR was calculated as the average HR before (ECG1) and after (ECG2) each sensory test (see Figure 1B).

#### 2.4.3. Glucose

Glucose concentrations were recorded using a continuous glucose monitoring (CGM) system (FreeStyle Libre 2). The CGM sensor was placed on the back of the upper arm on day one and glucose levels were obtained every 15 minutes throughout the five-days study. Glucose values were subsequently interpolated to 1-min intervals using MATLAB (MathWorks, Natick, MA, USA).

#### 2.4.4. Sleep

Sleep polysomnography was recorded every night from habitual bedtime to habitual wake time using a Vitaport-4 digital recorder (TEMEC Instruments, Kerkrade, Netherlands). Four electroencephalogram (EEG) electrodes were placed on the scalp at the C3, C4, O1, and O2 sites, according to the 10-20 international system. They were referenced to the contralateral mastoids (A1 and A2). Two electro-oculogram (EOG) electrodes were placed at 1 cm from the external canthi of the eyes and cross-referenced. Three electromyogram (EMG) electrodes were placed on the chin muscle (upper right, upper left, and lower center) and referenced (left-center and right-center). All signals were filtered using a high- and low-pass filter at 0.3 and 45 Hz, respectively, as well as a notch filter at 50 Hz. The sampling rate was 256 Hz.

### 2.5. Statistical analysis

Statistical analyses were conducted using RStudio (version 2023.06.2, PBC, Boston, MA, USA). All analyses (hunger, satiety, heart rate, blood glucose, total sleep time, and sleep latency (SL) were based on raw data. To eliminate the “first day effect,” analyses were performed from Tuesday (day 2, D2) to Friday (day 5, D5), with data collected before and after each of the three daily meals.

To ensure the reliability of the analyses, outliers were systematically identified and removed using a robust statistical approach. Specifically, outliers were detected using the interquartile range (IQR) method, in which outliers were defined as data points falling outside the acceptable range: below the first quartile (Q1 - 1.5 IQR) or above the third quartile (Q3 + 1.5 IQR).

Pre- and post-hunger/satiety were defined as the mean of the two measures around a sensory test. For example, pre-hunger is the mean of H1 and H2 around the pre-meal sensory test (see Figure 1B).

To examine how meal type (B, L, D) or day (D2 to D5), affected interoception, we applied a linear mixed-effects model that considered meal type, and the interaction between day and meal type. When the interaction was significant, we examined the simple effects (the effect of meal type on a particular day or the effect of day on a particular meal type). When there was no interaction, we considered a one-factor model.

To investigate whether different meals (breakfast, lunch, and dinner) influence interoception (hunger and satiety) during a day, we analyzed hunger and satiety separately under pre- and post-meal conditions separately. Then, we applied a linear mixed-effects model to the data. The model specification for pre-meal hunger, for example, was as follows: *PreHunger <-lme (Hunger ∼ Meal type + Day + Meal type x Day, random = 1 | Participants)*. When no significant interaction was found, we focused on the main effect (meal type or day). When an interaction was statistically significant, simple effects were tested. This model analyzed how pre-meal hunger perception varied as a function of meal type. The random factor, participants, was included to account for individual variability due to uncontrolled factors (Cheon & Mattes, 2024; Ruddick-Collins et al., 2019; Stevenson et al., 2015). Post hoc analysis with Bonferroni correction was used to adjust for multiple comparisons. The same analyses were performed for post-meal hunger, and pre- and post-meal satiety. Similar models were applied to physiological markers such as heart rate (HR) and glucose levels separately before and after meal.

To examine how individual interoception varies from day to day, we compared the within-subject variability over four days, and the between-subject variability for each type of meal (F-tests). This analysis quantifies how much of the total variation in interoception is attributable to intra-individual versus inter-individual differences. We applied the same approach to post-meal hunger and pre- and post-meal satiety, and physiological measures, including heart rate and glucose levels.

We applied a linear mixed model to assess the effect of low artificial light at night on sleep characterization indicators such as total sleep time (TST) and sleep latency (SL). The example model as follows: *TST <-lme (Total sleep time ∼ Light, random= ∼1 | Participant)*. We used a similar approach to analyze SL.

To investigate whether low artificial light at night influences the following day’s measured interoception, we applied a linear mixed model respectively for each meal (B, L, D) and each interoception condition (pre- or post-meal). *PreHungerBreakfast <-lme (Hunger ∼ Light, random= ∼1 | Participant)*. Similar models were applied to post-hunger, and pre- and post-satiety conditions.

To better understand the relationships between interoception (e.g., hunger, satiety) and physiological parameters (HR, glucose), Pearson correlation coefficients (*r*) were calculated 1) on pre- and post-meal data pooled together, 2) on pre-meal data, and 3) on post-meal data. Analyses were investigated in function of meal conditions (B, L, D). Pearson correlation analyses were also performed between sleep indicators such as total sleep time (TST) and sleep latency (SL), and the following day’s interoception measures.

All findings were considered statistically significant when *p* <= 0.05. Data are presented as the mean ± standard error of the mean (SEM) in the graphs.

## 3. Results

The framework of our highly controlled laboratory study conducted under time-isolation conditions, is shown in Figure 2 for interoception: within-subject longitudinal monitoring, three meals per day (averaged here), four consecutive days. The analyses on physiological data was computed using the same methodological framework and analytical procedure.

**Figure 2.**
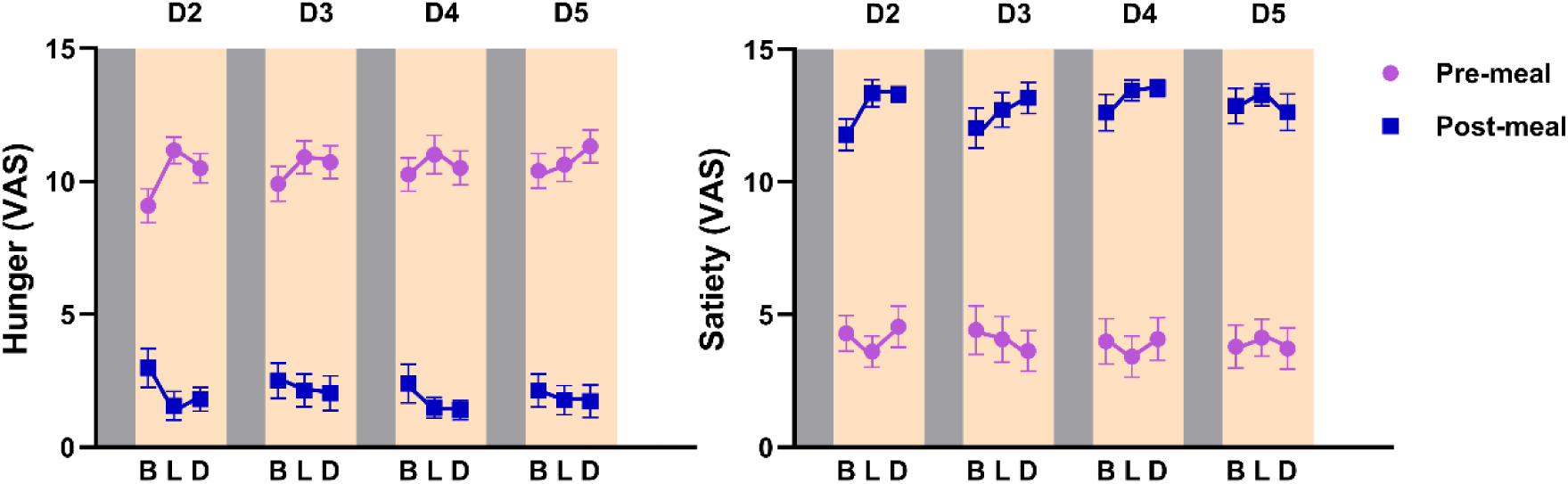
Hunger (left) and satiety (right) levels at pre- and post-meals in function of meal type and over four consecutive days D2 (Day 2) to D5 (Day 5). Data presented are means ± SEMs.

### 3.1. Interoception

#### 3.1.1. Fluctuation during the day

In this study, although both hunger and satiety were measured before and after each meal. In real-life conditions, these two indicators of interoception occur only under different metabolic states. Hunger occurs typically before meals, while satiety after meals. Therefore, the following results will only consider hunger before meals and satiety after meals. Since the preliminary analysis showed no interaction between days and meal types, the meal type effect reported here is the mean of all days.

The linear mixed-effects model revealed a significant main effect of meal type on hunger levels (*F*(2, 206) = 8.43, *p* < 0.0001). Post hoc comparisons showed that hunger levels before meals were significantly higher at lunch (11.25 ± 0.4, *p* < 0.001) and dinner (11.19 ± 0.4, *p* < 0.001) compared to breakfast (10.37 ± 0.4), with no significant difference between lunch and dinner (see Figure 3A).

**Figure 3.**
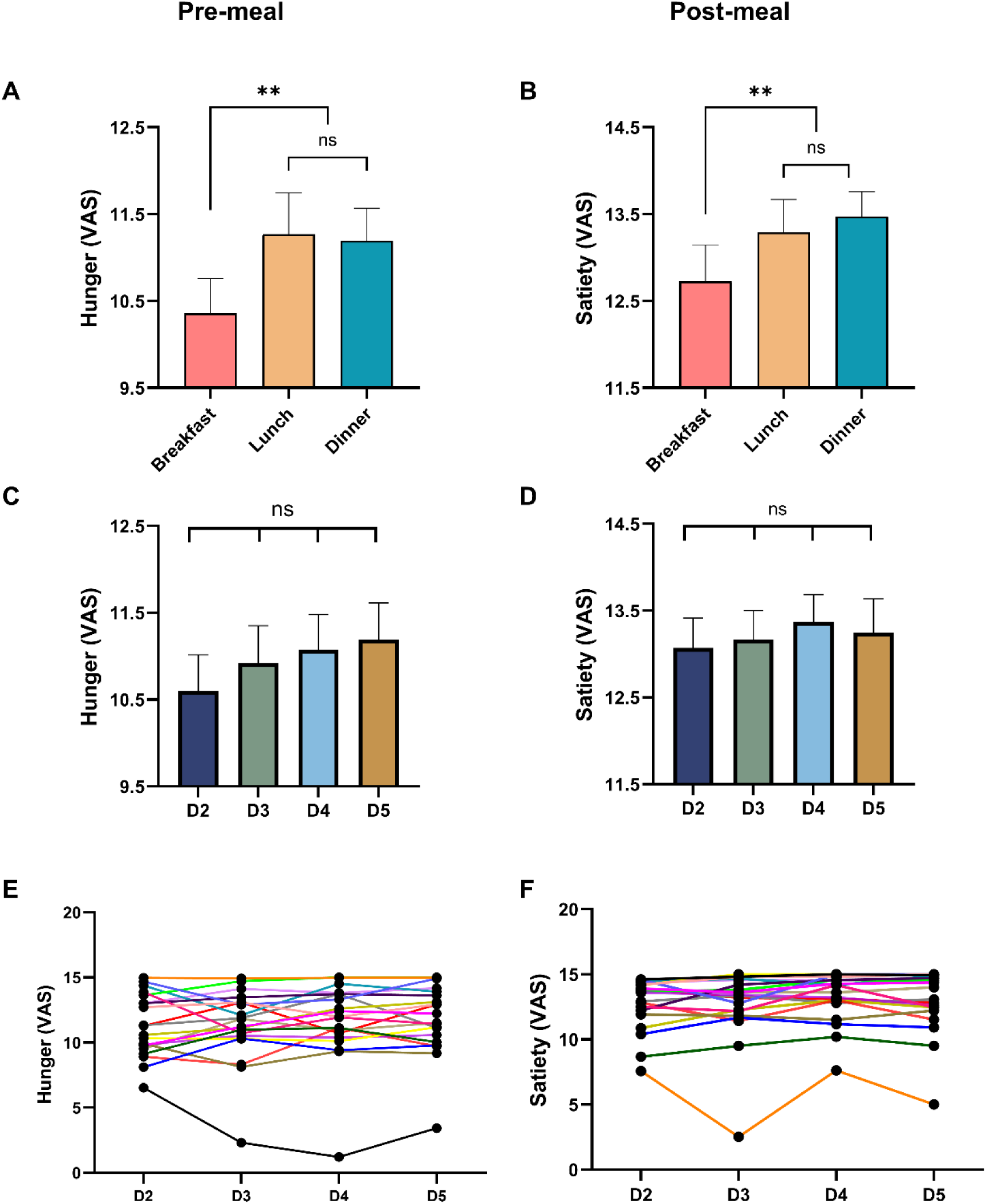
Dynamics and stability of hunger and satiety rating across the study period. Panels **(A-B)** illustrate the dynamics of hunger ratings measured before meal (A) and satiety rating measured after meals (B) at breakfast, lunch, and dinner. Panels (**C-D)** show the group mean hunger measured before meal (C) and satiety measured after meal (D) day 2 (D2) to day 5 (D5). Panels **(E-F)** show each subject’s individual hunger (E) and satiety (F) from D2 to D5. Each point represents the average values across the three meals (breakfast, lunch, dinner), and each colored line corresponds to the raw data from an individual participant. Data on histograms (A-D) are presented as means ± SEM.

The same model was applied to satiety (after meals), and showed a significant main effect of meal type throughout the day (*F*(2, 208) = 7.56, *p* < 0.0001). Post hoc comparisons revealed that satiety was significantly higher at lunch (13.4 ± 0.34, *p* < 0.001) and dinner (13.5 ± 0.34, *p* < 0.001) than at breakfast (12.93 ± 0.34), with no significant difference between lunch and dinner (see Figure 3B).

#### 3.1.2. Stability throughout the week

To examine how hunger and satiety evolved throughout the week, we first assessed mean values from D2 to D5. Hunger levels remained stable during experience (Figure 3C, D2: 10.6 ± 0.41; D3: 10.9 ± 0.41; D4: 11.1 ± 0.41; D5: 11.2 ± 0.41). A similar pattern was observed for satiety levels (Figure 3D, D2: 13.1 ± 0.35; D3: 13 ± 0.35; D4: 13.4 ± 0.35; D5: 13.3 ± 0.35). This stability across days was also evident when the three meals were analyzed separately. To further investigate day-to-day variations at the individual level, we compared within-individual variability (over four days) with between-individual variability (cumulated over the four days). We found that between-individual variability was significantly greater than within-individual variability for both hunger (Figure 3E, *F*(76, 60) = 10.5, *p* < 0.0001) and satiety (Figure 3F, *F*(76, 60) = 9.37, *p* < 0.0001).

### 3.3. Physiological parameters

#### 3.3.1. Heart rate

##### Pre-meal heart rate

Significant main effects of meal type (*F*(2, 208) = 18.86, *p* < 0.0001) and day (*F*(3, 208) = 3.03, *p* = 0.03) were observed for pre-meal Heart rate (HR), with no significant interaction between them (*F*(6, 208) = 1.3, *p* = 0.26). Across all day, pre-meal HR was significantly higher at B (66.4 ± 1.86 bpm) than at L (63.3 ± 1.86 bpm, *p* < 0.0001) and D (62.9 ± 1.86 bpm, *p* < 0.0001). There was no significant difference between L and D (*p* = 0.99) (see Figure 4A). Although a significant main effect of day was detected, post hoc analyses revealed only one significant difference: pre-meal HR was lower on D2 (63.2 ± 1.88 bpm) than on D5 (65.1 ± 1.88 bpm, *p* = 0.04; see Figure 4B).

**Figure 4.**
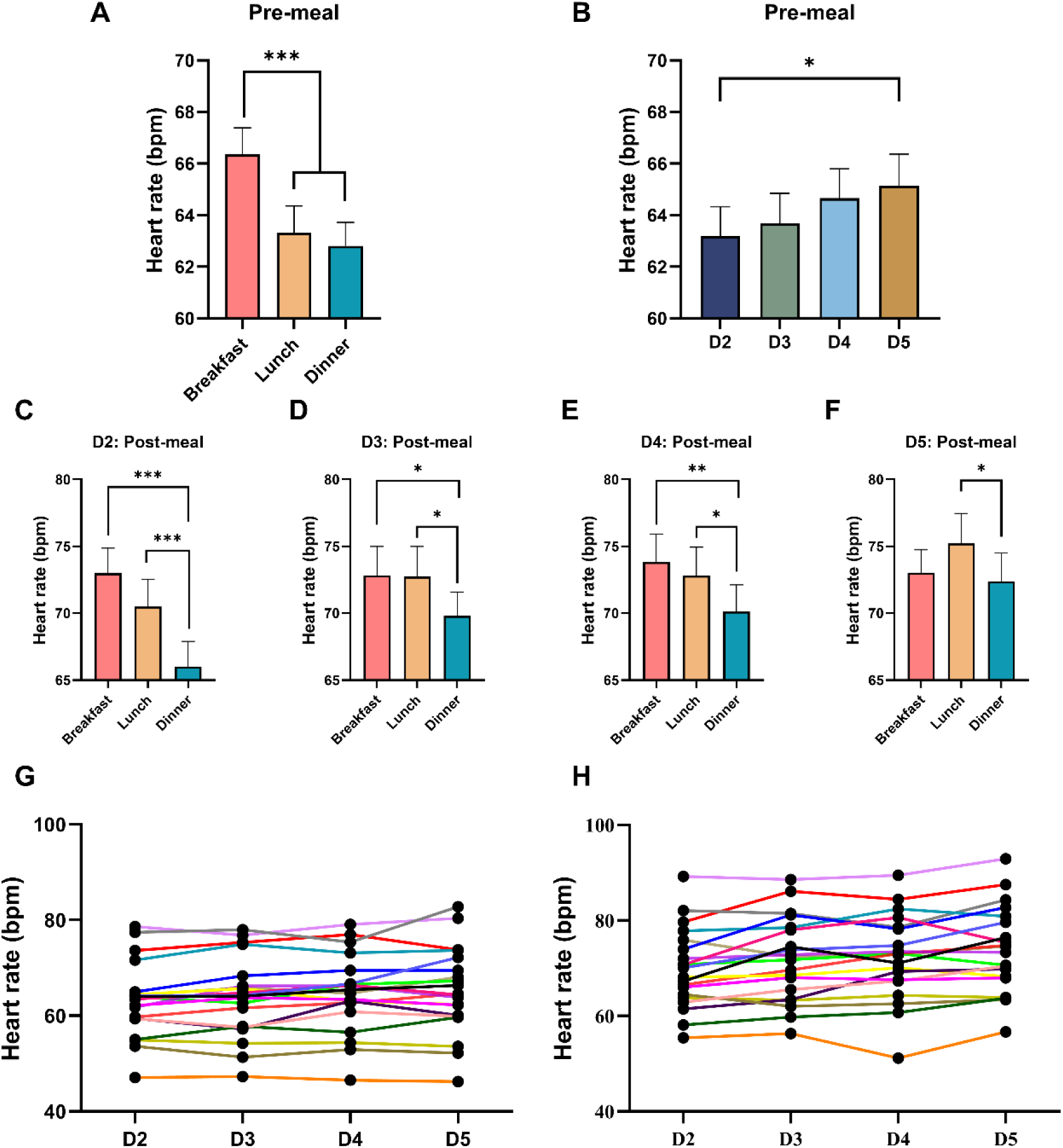
Heart rate (HR) before and after meals and across the study period. Panel **(A)** shows pre-meal Heart rate in function of meal types (breakfast, lunch, dinner). Panel **(B)** shows the group mean pre-meal HR from day 2 (D2) to day 5(D5). Panels (**C-F**) illustrate the dynamics of post-meal HR according to meal type on each study day: D2 (C), D3 (D), D4 (E), D5 (F). Panels (**G-H**) each subject’s individual pre-meal HR (G) and post-meal HR (H) from D2 to D5. Each point represents the average values across the three meals (Breakfast, lunch, and dinner), and each colored line corresponds to the raw data of an individual participant. Data on histograms (A-F) are presented as means ± SEM.

##### Post-meal heart rate

Significant main effects of meal type (*F*(2, 208) = 27.55, *p* < 0.0001) and day (*F*(3, 208) = 13.45, *p* < 0.0001), as well as a significant meal type x day interaction (*F*(6, 208) = 2.29, *p* = 0.037) on HR. The significant meal x day interaction comes from the fact that HR was significantly higher after breakfast (73.2 ± 2.05) than after dinner (68.63 ± 2.05), but only on days 2, 3, and 4 (D2, D3 and D4) (*p* < 0.05; Figures 4C-E). There was no such difference on day 5 (D5) (72.4 ± 2.05, *p* = 0.5; see Figure 4F). In addition, as shown in Figures 4C-F, HR was significantly higher after lunch than after dinner (69.58 ± 2.05) on all days.

##### Effect of meal consumption

Comparison of HR between pre- and post-meal revealed a significant increase in HR from pre- to post-meal for all meal types. HR increased respectively from 66.20 to 73.17 bpm for breakfast (*t* = 4.96, *p* < 0.0001), from 63.32 to 72.82 bpm for lunch (*t* = 6.34, *p* < 0.0001), and from 62.89 to 69.57 bpm for dinner (*t* = 4.96, *p* < 0.0001).

##### Inter-individual variability

At the individual level, between-individual variability was significantly greater than within-individual variability for both pre-meal HR (*F*(76, 60) = 31.73, *p* < 0.0001; Figure 4G) and post-meal HR (*F*(76, 60) = 16.25, *p* < 0.0001; Figure 4H).

#### 3.3.2. Glucose

##### Pre-meal glucose

Pre-meal blood sugar levels remained stable across the three meals, with no significant main effect of meal type (*F*(2, 205) = 2.21, *p* = 0.11; see Figure 5A). Likewise, there was no significant interaction between meal type and day (*F*(6, 205) = 1.59, *p* = 0.15).

**Figure 5.**
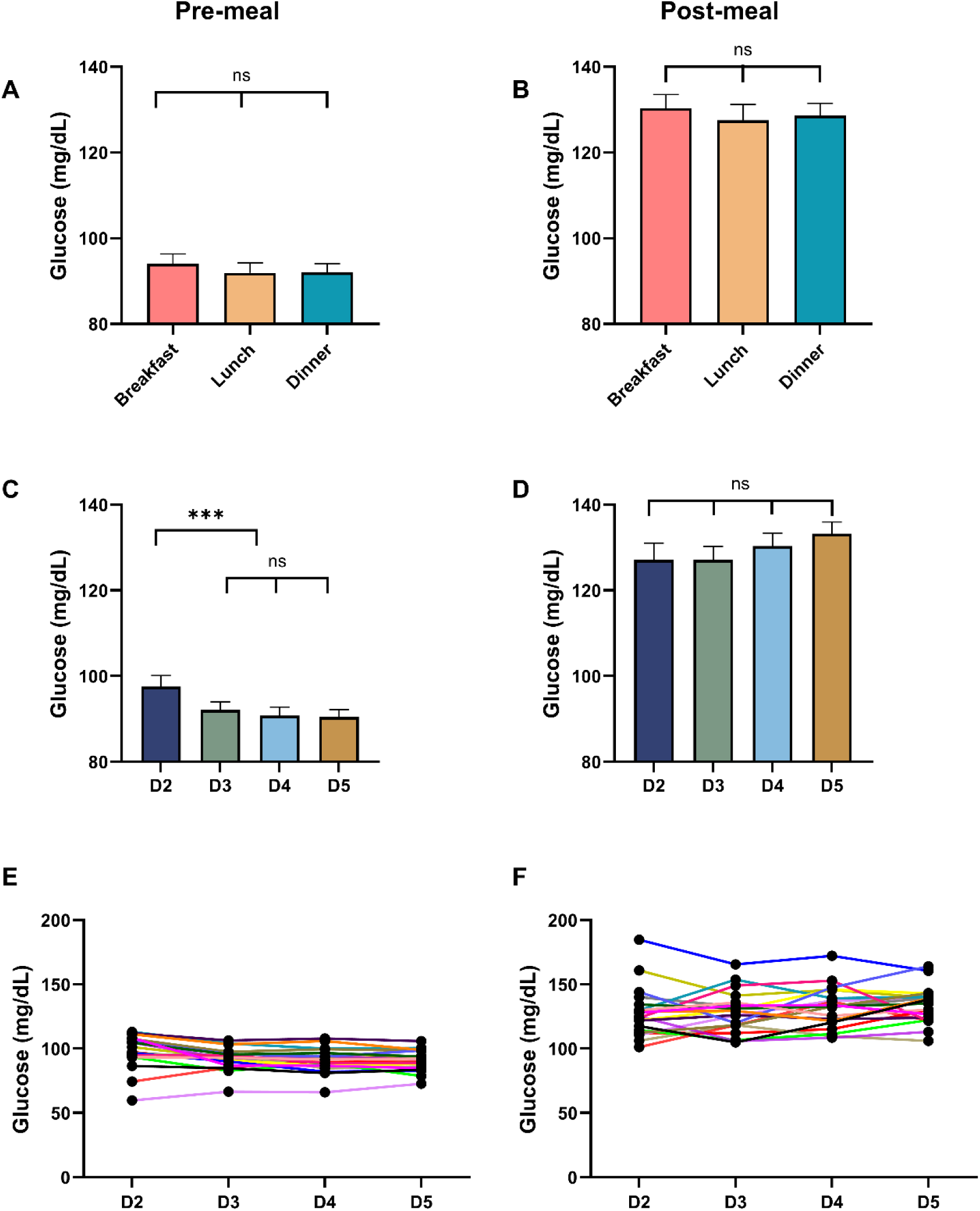
Glucose levels before and after meals and across the study period. Panels **(A-B)** illustrate glucose levels measured before meals (A) and after meals (B) at breakfast, lunch, and dinner. Panels (**C-D)** show the group mean glucose levels measured before meals (C) and after meals (D) from day 2 (D2) to day 5(D5). A significant effect of study day was observed for pre-meal glucose levels, with values on D2 differing significantly from those on the other study days. Panels **(E-F)** show each subject’s individual glucose level, measured before-meal (E) and after-meal (F), from D2 to D5. Each point represents the average values across the three meals (breakfast, lunch, dinner), and each colored line corresponds to the raw data from an individual participant. Data on histograms (A-D) are presented as means ± SEM.

##### Post-meal glucose

Post-meal blood sugar levels also remained stable across the three meal types, with no significant main effect of meal type (*F*(2, 203) = 1.3, *p* = 0.27; see Figure 5B). Similarly, there was no significant interaction between meal type and day (*F*(6, 203) = 0.34, *p* = 0.92).

##### Effect of study day

A significant main effect of study day was observed for pre-meal glucose levels (*F*(3, 205) = 15.39, *p* < 0.0001). Post hoc comparisons indicated that glucose concentrations on D2 (97.9 ± 2.0 mg/dL) were significantly higher than those on D3 (92.2 ± 2.0 mg/dL), D4 (90.8 ± 2.0 mg/dL), and D5 (90.4 ± 2.0 mg/dL; all *p* < 0.0001; see Figure 5C). In contrast, post-meal glucose levels did not differ significantly across study days (*F*(3, 203) = 2.51, *p* = 0.06; see Figure 5D).

##### Inter-individual variability

At the individual level, between-individual variability was significantly greater than within-individual variability for both pre-meal glucose levels (*F*(76, 60) = 5.3, *p* < 0.0001; Figure 5E) and post-meal glucose levels (*F*(76, 60) = 4.13, *p* < 0.0001; Figure 5F).

### 3.4. Correlations between hunger, satiety and physiological parameters

#### 3.4.1. Interoception: hunger and satiety

Hunger and satiety were negatively correlated in the pre- (*r*(226) = -0.68, *p* < 0.0001) and post-meal(*r*(219) = -0.72, *p* < 0.0001) states, respectively. In addition, a significant positive correlation was found between pre-meal hunger and post-meal satiety (*r*(216) = 0.3, *p* < 0.0001).

#### 3.4.2. Interoception and heart rate (HR)

When pre- and post-meal data pooled, a significant negative correlation was observed between hunger and HR on for all three meals combined (*r*(474) = -0.42, *p* < 0.0001), as well as three meals analyzed separately: breakfast (*r*(157) = -0.42, *p* < 0.0001), lunch (*r*(156) = - 0.47, *p* < 0.0001), and dinner (*r*(157) = -0.39, *p* < 0.0001; see Figure 6A). We also found a positive correlation between satiety and HR when the three meals were analyzed together (*r*(474) = 0.40, *p* < 0.0001) or separately: breakfast (*r*(157) = 0.42, *p* < 0.0001), lunch (*r*(156) = 0.45, *p* < 0.0001), and dinner (*r*(157) = 0.35, *p* < 0.0001; see Figure 6B).

**Figure 6.**
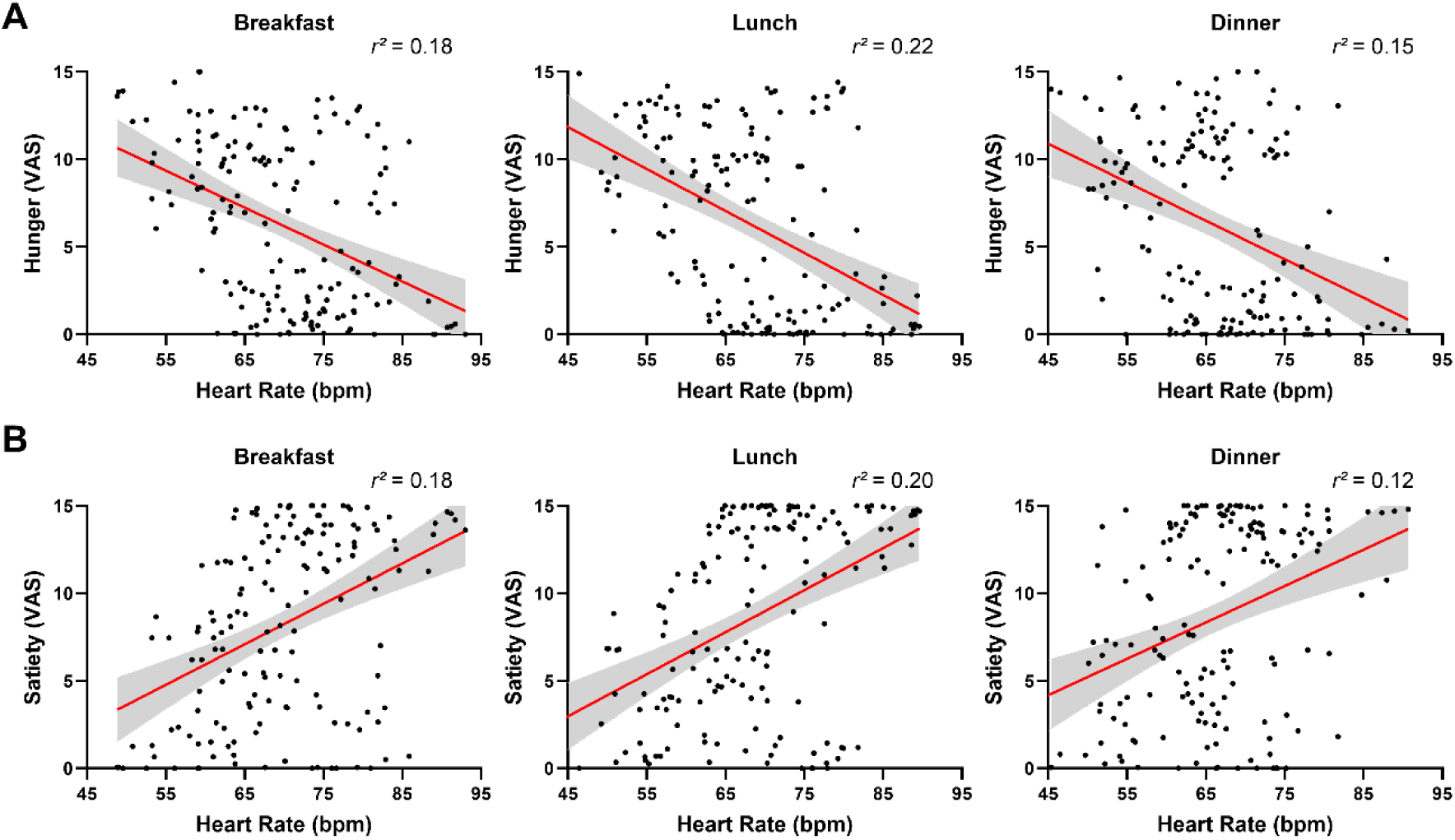
Association between interoception and HR when pre- and post-meal data were pooled. **(A)** Correlations between hunger ratings and HR at breakfast, lunch, and dinner. Significant negative correlations were observed for all meals (all p < 0.001). **(B)** Correlations between satiety ratings and HR at breakfast, lunch and dinner. Significant positive correlations were observed for all meals (all p < 0. 001).

When pre- and post-meal data were analyzed separately, a significant positive correlation was found only between post-meal satiety and HR (*r*(227) = 0.25, *p* = 0.0001). When meals analyzed individually, for pre-meal: no significant correlation was found between hunger and HR at breakfast (*r*(75) = -0.03, *p* = 0.75), lunch (*r*(77) = -0.028, *p* = 0.81), dinner (*r*(76) = -0.05, *p* = 0.64). In contrast for post-meal, significant positive correlations were observed between satiety and HR at breakfast (*r*(74) = 0.30, *p* = 0.009) and lunch (*r*(78) = 0.38, *p* = 0.0007), but no significant correlation at dinner (*r*(77) = 0.14, *p* = 0.24).

#### 3.4.3. Interoception and glucose levels

When pre- and post-meal data were pooled, a significant negative correlation was observed between hunger and glucose either when all three meals were combined (*r*(473) = -0.61, *p* < 0.001) or when analyzed separately (breakfast: (*r*(154) = -0.62, *p* < 0.0001; lunch: (*r*(158) = - 0.64, *p* < 0.0001; dinner: (*r*(157) = -0.58, *p* < 0.001; see Figure 7A). On the opposite, a significant positive correlation was observed between satiety and glucose levels, both when the three meals were combined (*r*(473) = 0.62, *p* < 0.001) and when analyzed separately (breakfast: (*r*(154) = 0.65, *p* < 0.0001; lunch: (*r*(158) = 0.62, *p* < 0.0001; dinner: (*r*(157) = 0.60, *p* < 0.0001; see Figure 7B).

**Figure 7.**
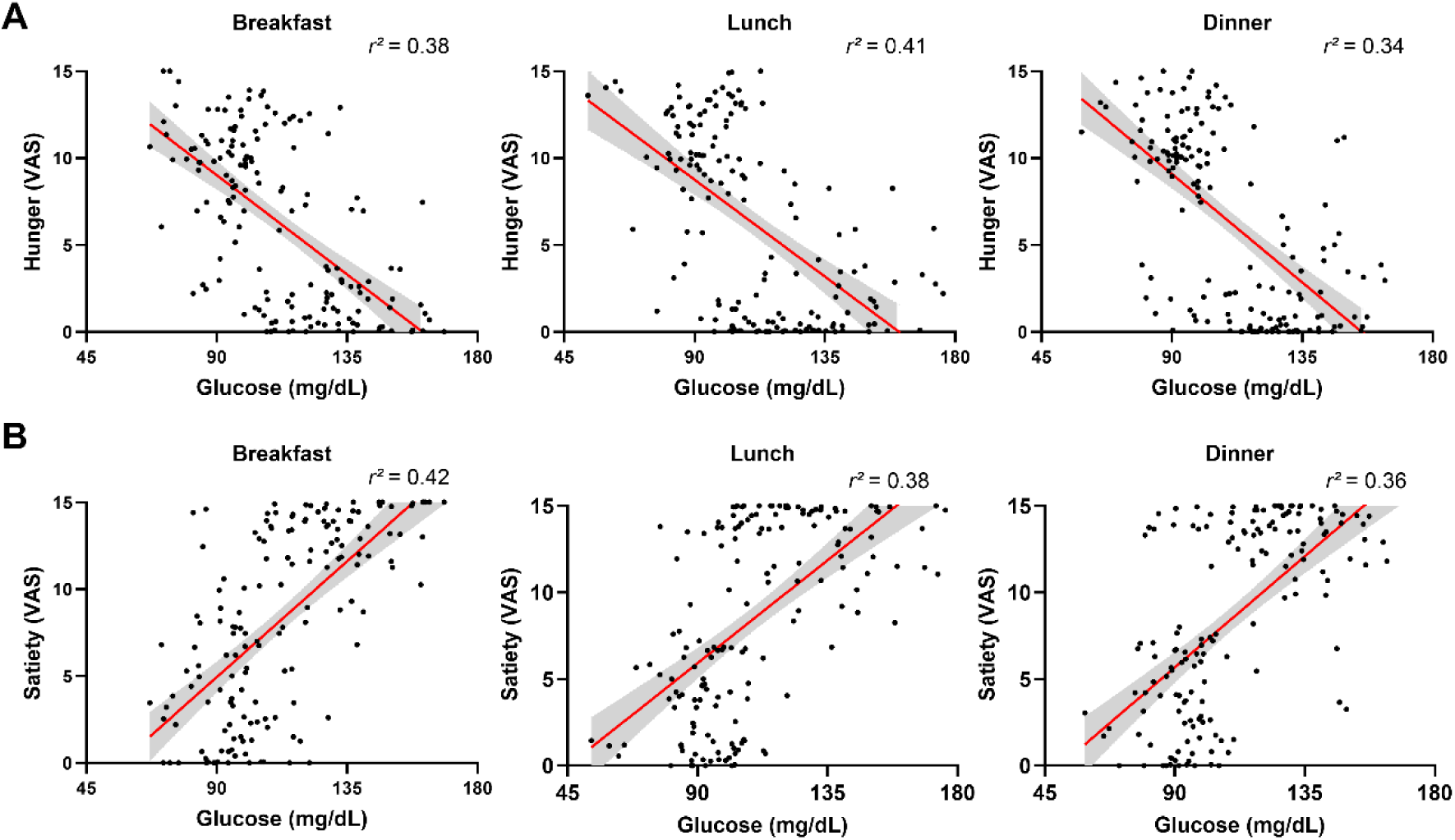
Association between interoception and glucose levels when pre- and post-meal data were pooled. **(A)** Correlations between hunger ratings and glucose levels at breakfast, lunch, and dinner. Significant negative correlations were observed for all meals (all p < 0.001). **(B)** Correlations between satiety ratings and glucose levels at breakfast, lunch and dinner. Significant positive correlations were observed for all meals. (all p < 0.001).

When pre- and post-meal data were analyzed separately, no significant correlations were observed between glucose levels and pre-meal hunger ratings (*r*(222) = -0.02, *p* = 0.80) or post-meal satiety (*r*(222) = -0.08, *p* = 0.23) when the three meals of the day were combined. Similarly, when meals were analyzed individually, no significant correlations were found between glucose levels and pre-meal hunger (breakfast: (*r*(75) = -0.06, *p* = 0.61; lunch: (*r*(74) = 0.17, *p* = 0.16; dinner: (*r*(75) = -0.13, *p* = 0.25) or post-meal satiety (breakfast: (*r*(71) = 0.19, *p* = 0.11; lunch: (*r*(78) = -0.21, *p* = 0.06; dinner: (*r*(75) = -0.16, *p* = 0.16).

### 3.5. Effects of low light intensity at night on interoception and sleep

No significant effect of light exposure during the night on pre-meal hunger of the following day was found when all three meals were combined (*F*(3, 205) = 0.80, *p* = 0.50) nor when the meals were analyzed separately: breakfast (*F*(3, 53) = 0.47, *p* = 0.70), lunch (*F*(3, 54) = 1.36, *p* = 0.26), and dinner (*F*(3, 54) = 1.36, *p* = 0.26). Similarly, no significant effect of light on post-meal satiety was found when all three meals were combined (*F*(3, 207) = 0.20, *p* = 0.89) nor when meals were analyzed separately: at breakfast (*F* (3, 52) = 1.06, *p* = 0.37), lunch (*F*(3, 55) = 0.94, *p* = 0.43) and dinner (*F*(3, 54) = 0.29, *p* = 0.83).

To investigate the potential indirect link between light exposure at night and the following day’s interoception, we analyzed the relationship between light exposure at night and sleep duration (total sleep time [TST]) and then TST and subsequent daytime interoception. No effect of light at night on total sleep time was found (*F*(3, 52) = 1.31, *p* = 0.28). However total sleep time (TST) was significantly, but weakly correlated with pre-meal hunger when all three meals were combined (*r*(216) = 0.24, *p* = 0.0004). When meals were analyzed separately, this correlation was significant for breakfast (*r*(70) = 0.24, *p* = 0.04) and lunch (*r*(71) = 0.33, *p* = 0.004), but not for dinner (*r*(71) = 0.13, *p* = 0.25). Furthermore, TST was not correlated with post-meal satiety, either when meals were combined (*r*(216) = 0.06, *p* = 0.38) or when examined separately for breakfast (*r*(69) = 0.01, *p* = 0.90), lunch (*r*(72) = 0.02, *p* = 0.89), and dinner (*r*(71) = 0.18, *p* = 0.16).

## 4. Discussion

The current study examined the characteristics of food-related interoception using a highly controlled, five-day laboratory protocol. Hunger and satiety were measured before and after each of 14 consecutive meals in 20 male subjects. Our results showed that interoception (hunger and satiety): 1) fluctuates throughout the day (daily variation); 2) remains stable across days; 3) is regulated by distinct mechanisms; 4) correlates with physiological parameters, such as heart rate and glycemia, and is partially correlated with prior sleep duration (pre-meal hunger).

### Dynamic interoception during the day

Hunger and satiety are momentary sensations that reflect respectively the body’s different metabolic and energy states (Schwartz et al., 2000; Woods et al., 1998), especially when the stomach is empty or full. Hunger typically increases before a meal and decreases and disappears with food intake, while satiety shows the opposite pattern. In this study, we found that these sensations vary depending on the meal type during a day. Specifically, pre-meal hunger and post-meal satiety are lower at breakfast than at lunch or dinner. These results suggest that hunger and satiety signals fluctuate throughout the day, with weaker perceptions in the morning (around breakfast) and stronger perceptions at midday (around lunch) and in the evening (around dinner).

Several studies have investigated interoception throughout the day, assessing hunger levels multiple times. These studies have suggested that hunger may fluctuate on a daily basis. A weak sense of hunger is typically observed at the start of the day and becomes progressively stronger hunger later in the day (Cugini et al., 1998; Fatati et al., 2001; Qian et al., 2019; Sargent et al., 2016; Scheer et al., 2013). However, fewer studies have examined variation in satiety throughout the day. One exception is the study by Sargent et al. (2016), which measured hunger and satiety in a 28-hour forced desynchrony protocol. While our results regarding dynamic changes in hunger throughout the day are similar to that of Sargent and colleagues, we find the opposite result for satiety. Indeed, contrary to us, Sargent et al. (2016) found the lowest satiety levels in the evening (5:00 p.m. to 9:00 p.m.) and the highest levels at night (1:00 a.m. to 5:00 a.m.). One possible explanation for this difference is that the authors did not specify when the satiety measurements were taken relative to meal times (i.e., before or after). If satiety and hunger were assessed simultaneously, the expected negative correlation between the two could result in an inverse relationship, whereby lower hunger would correspond to higher satiety. Another reason is that Sargent and his colleagues used a forced-desynchrony protocol, which may have resulted in progressive sleep debt accumulation throughout the study, and influenced the participants’ interoceptive dynamics.

The sustained higher pre-meal hunger and post-meal satiety experienced at lunch and dinner may have evolutionary significance. A strong hunger signal at the end of the day could increase motivation to eat. Concurrently, the ability to achieve greater satiety levels could extend the time period during which food is consumed. Together, these two mechanisms ensure adequate energy availability at the end of the day (lunch-related), and in anticipation of the overnight fasting sleep episode (diner-related). However, in today’s obesogenic environment, resulting from food availability and attractivity, and influenced by psychological factors (e.g. expectation and motivation) (de-Arruda et al., 2024; Mathew et al., 2022), such a dual mechanism may lead to overeating at lunch and dinner. The increasing tendency to skip breakfast and curtail our sleep (i.e., carry a sleep debt) in our modern societies may constitute an additional vulnerability leading to an increased food intake before bedtime, consequently contributing to an increased risk of cardiometabolic disorders (Gronfier et al., 2024).

### Stability of interoception from one day to another

Although interoception fluctuates throughout the day, our findings indicate that it remained stable over the course of the four-day study. Each participant exhibited consistent hunger and satiety levels respectively before or after each of the three meal types (B, L, D). Intra-individual variability over the four days was significantly lower than inter-individual variability. In other words, differences in interoception between individuals were far greater than the day-to-day fluctuations observed within the same individual.

While several studies have measured hunger or fullness (satiety) over multiple days or weeks (Cheon & Mattes, 2024; Ruddick-Collins et al., 2019; Stevenson et al., 2015), few have specifically examined longitudinal changes in interoception over time. Only the study by Cheon and Mattes (2024) found that hunger and satiety remained relatively stable when comparing measures taken from the same subjects three times, with eight weeks apart (weeks 1, 9, and 17). They also reported considerable interindividual variability, with which our findings are consistent.

The stability of individual interoception observed in our study is unlikely to be due to the habitual use of the line scale or repeated measures of interoception. Participants reported lower levels of hunger and satiety in the morning, with higher ratings at lunch and dinner. Since hunger and satiety are regulated by stomach fullness and hormones such as ghrelin and leptin (Cummings et al., 2001; Klok et al., 2007), the consistency of interoception across days may rather reflect stable individual hormone secretion patterns (Scheer et al., 2009) or metabolic states at specific times. This stability may also be linked to a stable sleep-wake cycle and circadian rhythmicity, which are inherent to our highly controlled pre-laboratory and laboratory conditions.

These findings suggest that the daily fluctuations in hunger and satiety, as well as stability over times, represent characteristic individual traits. The level of an individual’s interoception may be used to differentiate subjects. This individual trait, the perception of hunger and satiety, may result from long-term physiological adaptations that may contribute to the maintaining stable food intake and body weight over time.

### Hunger and satiety are closely related but two distinct parameters

Hunger and satiety are generally considered to be two opposing interoceptive states that reflect energy deficits and surpluses, respectively. Consequently, most studies usually assess only one of them. However, when both are measured simultaneously, they are typically negatively correlated (Scheer et al., 2013). But is it necessary and pertinent to measure both simultaneously? Although hunger and satiety are statistically significantly correlated when measured simultaneously (*r* = -0.68 in our study), less than 50% of the variation of hunger can be explained by satiety variation, Thus, the two are not perfectly opposite interoceptions. In fact, hunger and satiety operate under different physiological conditions and rely on different neural circuits and neurotransmitters to control different steps of food intake (Amin & Mercer, 2016; Campfield & Smith, 1990b, 1990a; Cummings, 2006; Cummings et al., 2001; Schwartz et al., 2000). Although both hunger and satiety are physiological and psychological sensations influenced by internal and external factors, they reflect different internal states: if one (e.g., hunger) is perceived, the other (satiety) is usually absent, and vice versa. Therefore, it is pertinent to measure hunger before food intake and satiety after. When we correlated the pre-meal hunger with post-meal satiety, the correlation was statistically significant, with a R^2^ at 0.09. This very small coefficient of determination favors our hypothesis that the two interoceptions function independently of each other, controlling different aspects of food intake (e.g., the motivation to initiate food intake and tolerance of ingestion quantity). This hypothesis is consistent with fingdings from two previous studies. Pélissier et al. (2024) reported that physical exercise can modify post-meal satiety perception without affecting pre-meal appetite-related measures. In addition, Fetissove et al. (2000) found that monoamine release in the medial and lateral hypothalamus was synchronized specifically during the satiety period. Therefore, since these two interoception are regulated differently, evaluating both is relevant and necessary to thoroughly examine their role in food intake regulation (Amin & Mercer, 2016; Nakamura & Nakamura, 2018). Furthermore, given the higher signal to noise ratios observed for hunger rating before meals and satiety ratings after meals, we propose that hunger should be measured before and satiety after food intake in order to properly evaluate interoception dynamics and helpful to understand the between individual difference when using interoception in food intake control.

### Interoception and physiological parameters

The relationship between interoception and physiological parameters, such as heart rate and glycemia, has rarely been studied together. However, these parameters reflect different aspects of the internal states (Harthoorn & Dransfield, 2008).

### Heart rate and hunger/satiety

One finding of the present study is that heart rate is higher in a satiety state than in a hunger state for all meals. At first glance, this finding seemed contrary to the classical assumption. It is well known that the sympathetic system increases heart rate, while the parasympathetic activity decreases it. Since hunger is usually considered an internal signal that drives exploration and intake behavior, and satiety occurs during digestion, one might expect a satiety state to be linked to parasympathetic activity and associated with a reduced heart rate. The sympathetic system activates “fight or flight” responses, while the parasympathetic system predominates during “rest and digest”. Several studies have examined the relationship between metabolic state and cardiac activity under different nutritional conditions or at a particular metabolic state. However, none of these studies have provided evidence to support the hypothesis that heart rate shifts from hunger to satiety (Flasbeck et al., 2021; Herbert et al., 2012; Peschel et al., 2018; Polito et al., 2022). Nevertheless, our counterintuitive finding (higher heart rate at satiety) was supported by one study in which heart rate was recorded four times per hour during about five hours, covering a lunch period states (Harthoorn & Dransfield, 2008). This study showed that heart rate is higher after lunch. However, heart rate was not measured around the other meals, or over several days. Thanks to our design protocol, which included 14 consecutive measurements across five days, our study reveals that heart rate is higher in the satiety state after all 14 meals. Although few studies have directly measured the effects of meal on heart rate, several indirect studies support these results, as there is higher sympathetic tone associated with satiety (Harthoorn & Dransfield, 2008). The higher sympathetic activity after meals may be linked to the physiological regulation of blood flow redistribution in the body, although the precise mechanisms are still unknown. In addition, our study and the study by Harthoorn and Dransfield (2008) demonstrate that the increased higher heart rate following a meal persists for at least one hour. Therefore, altogether, these results may suggest a perspective hypothesis to verify: whether people who regularly eat until full satiety are at a higher risk for cardio-metabolic diseases due to their consistently higher HR after eating.

### Glycemia and hunger/satiety

Our results revealed modest correlations between glycemia and interoception when pre- and post-meal data were analyzed together. Increasing glucose concentrations were associated with reduced hunger and increased satiety, which is consistent with previous findings (Campfield et al., 1996; Melanson et al., 1999). Since hunger and satiety are recognized as physiological markers of energy status and involved in feeding control, these results suggest that glycemic signals play a role in appetite modulation. However, causality cannot be inferred from these present correlation analyses alone. Importantly, no significant associations were observed when pre- and post-meal conditions were analyzed separately. This suggests that the relationships identified in the pooled analysis primarily reflect the physiological transition between fasting and postprandial states rather than direct moment-to-moment coupling within each condition. These results further indicate that interoception is influenced by multiple mechanisms beyond glycemia, including gastrointestinal peptide signaling, gastric distension (Cummings & Overduin, 2007), sensory reward systems (Monteleone et al., 2012), and cognitive, behavioral factors (Berthoud, 2011).

### Interoception and sleep

We did not find a strong overall relationship between interoception and sleep, but only a positive correlation between pre-meal hunger (at breakfast and lunch) and sleep duration. This may be due to the fact that in the present study, light intensity was maintained at very low levels (0, 3, 8, and 20 lux) at night. These levels were selected to be compatible with real-life nighttime lighting conditions observed at home, contrasting with the higher intensities typically used in other studies (Chang et al., 2012; Gronfier et al., 2004, 2007; Najjar et al., 2024; Rahman et al., 2019; Zeitzer et al., 2000). Since these low light intensities did not significantly influence total sleep time or sleep latency, they may not have been sufficient to disrupt the relationship between interoception and sleep in a detectable way. It is also plausible that the homeostatic regulations of both sleep and energy balance were robust enough to maintain stability despite these low and unimpactful light intensities. The positive correlation we found between total sleep time (TST) and pre-meal hunger at breakfast and lunch suggests that, under good sleep conditions, as opposed to sleep deprivation conditions often used in other studies (Spiegel et al., 1999), adequate sleep might play a role in optimizing energy balance and promoting metabolic health.

Several limitations should be noted. First, our study was conducted under highly controlled laboratory conditions. Although appetite and hunger in real-life settings may be modulated by other environmental factors such as exercise and light exposure, our results showing systematic diurnal variation and stability over several days advocate for the existence of internal regulatory mechanisms. Secondly, the meals provided to our subjects for breakfast, lunch and dinner were not identical. This may have contributed to some of the observed diurnal variability; however, our results are consistent with previous studies suggesting a possible circadian rhythm in hunger perception (Scheer et al., 2013), supporting the notion of a dynamic and endogenous rhythm in interoception. Finally, since the study was conducted exclusively with male participants, and there may be gender difference in interoception (Monrroy et al., 2019) further research with a more diverse population is needed to ensure the generalizability of our findings.

## 5. Conclusion

From the data presented, we conclude that: 1) hunger and satiety are the two interoceptive processes that may act in concert but are distinct; 2) perceived levels of hunger and satiety fluctuate during the day, with lower intensities at the beginning of the day and higher intensities around noon and dinner; 3) each individual has unique perceived levels of hunger and satiety that remain stable over relatively long periods of time; 4) both pre-meal hunger and post-meal satiety measures are relevant and necessary to explaining food intake strategies; 5) heart rate increases in the satiety state. The relationship between these interoceptive processes and physiological parameters is complex. An increase in post-meal heart rate may be a useful indicator for predicting certain food-intake-related illnesses, but further investigation is required.

## 6. Conclusion

### Data availability declaration

The data that support the findings of this study are available from the corresponding author upon reasonable request.

### CRediT authorship contribution statement

**XW:** Investigation, Data curation, Formal analysis, Visualization, Validation, Writing – original draft, Writing – review and editing. **ASP:** Project administration, Investigation, Data curation, Visualization, Validation, Writing – review and editing. **NT:** Investigation, Data curation, Formal analysis, Visualization, Validation, Writing – review and editing. **YH:** Investigation, Data curation, Formal analysis, Visualization, Validation, Writing – review and editing. **MT**: Formal analysis, Resources, Software, Validation, Writing – review and editing. **CG:** Conceptualization, Methodology, Funding acquisition, Project administration, Investigation, Supervision, Data curation, Visualization, Validation, Writing – original draft, Writing – review and editing. **TJ:** Conceptualization, Methodology, Funding acquisition, Project administration, Investigation, Supervision, Data curation, Visualization, Validation, Writing – original draft, Writing – review and editing.

### Generative artificial intelligence usage

No generative AI tools were used in the preparation of this manuscript.

### Funding sources

This work was supported by grants from the ANR (Idex Breakthough ALAN, 16 IDEX-0005) and the University of Lyon (Rectolux) to CG, as well as by funds from Inserm and CNRS to CG and TJ.

## Acknowledgments

The authors would like to thank Anissa Dahmani and Lydie Merle for their contributions to this study. Finally, the authors thank all of the participants who were contributed to or were interested in this study.

## Ethics statement

All experimental procedures adhered to the Declaration of Helsinki, and the protocol was approved by the Institutional Review Board (Comité de Protection des Personnes Ouest VI, Brest, France, N°ID-RCB: 2023-A01561-44). Informed consent was obtained from all participants.

## References

1. Amin, T., & Mercer, J. G. (2016). Hunger and Satiety Mechanisms and Their Potential Exploitation in the Regulation of Food Intake. Current Obesity Reports, 5, 106–112. 10.1007/s13679-015-0184-5

2. Barbosa, D. E. C., Souza, V. R. de, Santos, L. A. S. dos, Chiappini, C. C. de J., Sa, S. A. de, & Azeredo, V. B. de. (n.d.). Changes in Taste and Food Intake during the Menstrual Cycle. Journal of Nutrition & Food Sciences, 5(4), 1–6. 10.4172/2155-9600.1000383

3. Beck, A. T., Ward, C. H., Mendelson, M., Mock, J., & Erbaugh, J. (1961). An inventory for measuring depression. Archives of General Psychiatry, 4, 561–571. 10.1001/archpsyc.1961.01710120031004

4. Bellisle, F. (2005). Faim et satiété, contrôle de la prise alimentaire. EMC - Endocrinologie, 2(4), 179–197. 10.1016/j.emcend.2005.08.003

5. Benton, M. J., Hutchins, A. M., & Dawes, J. J. (2020). Effect of menstrual cycle on resting metabolism: A systematic review and meta-analysis. PloS One, 15(7), e0236025. 10.1371/journal.pone.0236025

6. Berthoud, H.-R. (2011). Metabolic and hedonic drives in the neural control of appetite: Who’s the boss? Current Opinion in Neurobiology, 21(6), 888–896. 10.1016/j.conb.2011.09.004

7. Berthoud, H.-R., Morrison, C. D., & Münzberg, H. (2020). The obesity epidemic in the face of homeostatic body weight regulation: What went wrong and how can it be fixed? Physiology & Behavior, 222, 112959. 10.1016/j.physbeh.2020.112959

8. Blundell, J., de Graaf, C., Hulshof, T., Jebb, S., Livingstone, B., Lluch, A., Mela, D., Salah, S., Schuring, E., van der Knaap, H., & Westerterp, M. (2010). APPETITE CONTROL: METHODOLOGICAL ASPECTS OF THE EVALUATION OF FOODS. Obesity Reviews : An Official Journal of the International Association for the Study of Obesity, 11(3), 251–270. 10.1111/j.1467-789X.2010.00714.x

9. Brondel, L., Romer, M. A., Nougues, P. M., Touyarou, P., & Davenne, D. (2010). Acute partial sleep deprivation increases food intake in healthy men123. The American Journal of Clinical Nutrition, 91(6), 1550–1559. 10.3945/ajcn.2009.28523

10. Broussard, J. L., Kilkus, J. M., Delebecque, F., Abraham, V., Day, A., Whitmore, H. R., & Tasali, E. (2016). Elevated ghrelin predicts food intake during experimental sleep restriction. Obesity (Silver Spring, Md.), 24(1), 132–138. 10.1002/oby.21321

11. Buysse, D. J., Reynolds, C. F., Monk, T. H., Berman, S. R., & Kupfer, D. J. (1989). The Pittsburgh sleep quality index: A new instrument for psychiatric practice and research. Psychiatry Research, 28(2), 193–213. 10.1016/0165-1781(89)90047-4

12. Campfield, L. A., Brandon, P., & Smith, F. J. (1985). On-line continuous measurement of blood glucose and meal pattern in free-feeding rats: The role of glucose in meal initiation. Brain Research Bulletin, 14(6), 605–616. 10.1016/0361-9230(85)90110-8

13. Campfield, L. A., & Smith, F. J. (1990a). Systemic factors in the control of food intake: Evidence for patterns as signals. In Neurobiology of food and fluid intake (pp. 183–206). Plenum Press. 10.1007/978-1-4613-0577-4_8

14. Campfield, L. A., & Smith, F. J. (1990b). Transient declines in blood glucose signal meal initiation. International Journal of Obesity, 14 Suppl 3, 15–31; discussion 31-4.

15. Campfield, L. A., Smith, F. J., Rosenbaum, M., & Hirsch, J. (1996). Human eating: Evidence for a physiological basis using a modified paradigm. Neuroscience and Biobehavioral Reviews, 20(1), 133–137. 10.1016/0149-7634(95)00043-e

16. Candan, E., Metin, Z. E., & Tengilimoglu-Metin, M. M. (2025). The role of premenstrual syndrome in hedonic hunger and food craving during the menstrual cycle. Journal of Nutritional Science, 14, e66. 10.1017/jns.2025.10038

17. Chang, A.-M., Santhi, N., St Hilaire, M., Gronfier, C., Bradstreet, D. S., Duffy, J. F., Lockley, S. W., Kronauer, R. E., & Czeisler, C. A. (2012). Human responses to bright light of different durations. The Journal of Physiology, 590(13), 3103–3112. 10.1113/jphysiol.2011.226555

18. Cheon, E., & Mattes, R. D. (2024). Interindividual variability in appetitive sensations and relationships between appetitive sensations and energy intake. International Journal of Obesity (2005), 48(4), 477–485. 10.1038/s41366-023-01436-9

19. Cugini, P., Ventura, M., Ceccotti, P., Cilli, M., Marcianò, F., Salandri, A., Di Marzo, A., Fontana, S., Pellegrino, A. M., Vacca, K., & Di Siena, G. (1998). Hunger sensation: A chronobiometric approach to its within-day and intra-day recursivity in anorexia nervosa restricting type. Eating and Weight Disorders: EWD, 3(3), 115–123. 10.1007/BF03339998

20. Cummings, D. E. (2006). Ghrelin and the short- and long-term regulation of appetite and body weight. Physiology & Behavior, Making Sense of Food, 89(1), 71–84. 10.1016/j.physbeh.2006.05.022

21. Cummings, D. E., & Overduin, J. (2007). Gastrointestinal regulation of food intake. Journal of Clinical Investigation, 117(1), 13–23. 10.1172/JCI30227

22. Cummings, D. E., Purnell, J. Q., Frayo, R. S., Schmidova, K., Wisse, B. E., & Weigle, D. S. (2001). A preprandial rise in plasma ghrelin levels suggests a role in meal initiation in humans. Diabetes, 50(8), 1714–1719. 10.2337/diabetes.50.8.1714

23. Daguet, I., Raverot, V., Bouhassira, D., & Gronfier, C. (2022). Circadian rhythmicity of pain sensitivity in humans. Brain: A Journal of Neurology, 145(9), 3225–3235. 10.1093/brain/awac147

24. de-Arruda, J. P., de-Souza, A. P. A., Pereira, L. P., Fonseca, L. B., Nogueira, P. S., Rodrigues, P. R. M., Muraro, A. P., & Ferreira, M. G. (2024). Short Sleep Duration and Skipping Main Meals among University Students. Sleep Science (Sao Paulo, Brazil), 17(4), e414–e421. 10.1055/s-0044-1782178

25. Fatati, G., Vendetli, A. L., Puxeddu, A., De Francesco, G. P., Coda, S., De Rosa, R., De Marco, E., De Laurentis, T., Fontana, S., & Cugini, P. (2001). Circadian rhythm of hunger sensation in obese patients: Effects of a short-term, moderately hypocaloric diet with a substitutive meal. Eating and Weight Disorders: EWD, 6(4), 214–219. 10.1007/BF03339745

26. Fetissov, S. O., Meguid, M. M., Chen, C., & Miyata, G. (2000). Synchronized release of dopamine and serotonin in the medial and lateral hypothalamus of rats. Neuroscience, 101(3), 657–663. 10.1016/S0306-4522(00)00374-2

27. Flasbeck, V., Bamberg, C., & Brüne, M. (2021). Short-Term Fasting and Ingestion of Caloric Drinks Affect Heartbeat-Evoked Potentials and Autonomic Nervous System Activity in Males. Frontiers in Neuroscience, 15, 622428. 10.3389/fnins.2021.622428

28. Garner, D. M., & Garfinkel, P. E. (1979). The Eating Attitudes Test: An index of the symptoms of anorexia nervosa. Psychological Medicine, 9(2), 273–279. 10.1017/s0033291700030762

29. Gorczyca, A. M., Sjaarda, L. A., Mitchell, E. M., Perkins, N. J., Schliep, K. C., Wactawski–Wende, J., & Mumford, S. L. (2016). Changes in macronutrient, micronutrient, and food group intakes throughout the menstrual cycle in healthy, premenopausal women. European Journal of Nutrition, 55(3), 1181–1188. 10.1007/s00394-015-0931-0

30. Green, S. M., Delargy, H. J., Joanes, D., & Blundell, J. E. (1997). A satiety quotient: A formulation to assess the satiating effect of food. Appetite, 29(3), 291–304. 10.1006/appe.1997.0096

31. Gronfier, C., Beaudart, C., Challet, E., Davenne, D., Delaunay, F., Duez, H., Galusca, B., Jacobi, D., Jiang, T., Morio-Liondore, B., Simonneaux, V., Mariotti, F., Barreau, F., Beaudart, C., Bennetau-Pelissero, C., Benzi-Schmid, C., Boutron-Ruault, M.-C., Lauzon-Guillain, B. D., Divaret-Chauveau, A., … Walrand, S. (2024). Actualisation des repères du PNNS: Répartition temporelle des prises alimentaires (Saisine n°2019-SA-0001; p. 202 p.). Anses. https://anses.hal.science/anses-04609761

32. Gronfier, C., Wright, K. P., Kronauer, R. E., & Czeisler, C. A. (2007). Entrainment of the human circadian pacemaker to longer-than-24-h days. Proceedings of the National Academy of Sciences, 104(21), 9081–9086. 10.1073/pnas.0702835104

33. Gronfier, C., Wright, K. P., Kronauer, R. E., Jewett, M. E., & Czeisler, C. A. (2004). Efficacy of a single sequence of intermittent bright light pulses for delaying circadian phase in humans. American Journal of Physiology - Endocrinology and Metabolism, 287(1), E174–E181. 10.1152/ajpendo.00385.2003

34. Harthoorn, L. F., & Dransfield, E. (2008). Periprandial changes of the sympathetic–parasympathetic balance related to perceived satiety in humans. European Journal of Applied Physiology, 102(5), 601–608. 10.1007/s00421-007-0622-5

35. Herbert, B. M., Muth, E. R., Pollatos, O., & Herbert, C. (2012). Interoception across modalities: On the relationship between cardiac awareness and the sensitivity for gastric functions. PloS One, 7(5), e36646. 10.1371/journal.pone.0036646

36. Horne, J. A., & Ostberg, O. (1976). A self-assessment questionnaire to determine morningness-eveningness in human circadian rhythms. International Journal of Chronobiology, 4(2), 97–110.

37. Jeon, B., & Baek, J. (2023). Menstrual disturbances and its association with sleep disturbances: A systematic review. BMC Women’s Health, 23(1), 470. 10.1186/s12905-023-02629-0

38. Jose, A. D., & Collison, D. (1970). The normal range and determinants of the intrinsic heart rate in man. Cardiovascular Research, 4(2), 160–167. 10.1093/cvr/4.2.160

39. Kabir, A., Miah, S., & Islam, A. (2018). Factors influencing eating behavior and dietary intake among resident students in a public university in Bangladesh: A qualitative study. PLoS ONE, 13(6), e0198801. 10.1371/journal.pone.0198801

40. Klok, M. D., Jakobsdottir, S., & Drent, M. L. (2007). The role of leptin and ghrelin in the regulation of food intake and body weight in humans: A review. Obesity Reviews: An Official Journal of the International Association for the Study of Obesity, 8(1), 21–34. 10.1111/j.1467-789X.2006.00270.x

41. Krishnan, S., Tryon, R., Welch, L. C., Horn, W. F., & Keim, N. L. (2016). Menstrual cycle hormones, food intake, and cravings. The FASEB Journal, 30(S1), 418.6-418.6. 10.1096/fasebj.30.1_supplement.418.6

42. Louis-Sylvestre, J. (1976). Preabsorptive insulin release and hypoglycemia in rats. The American Journal of Physiology, 230(1), 56–60. 10.1152/ajplegacy.1976.230.1.56

43. Marcelino, A. S., Adam, A. S., Couronne, T., Köster, E. P., & Sieffermann, J. M. (2001). Internal and external determinants of eating initiation in humans. Appetite, 36(1), 9–14. 10.1006/appe.2000.0375

44. Markwald, R. R., Melanson, E. L., Smith, M. R., Higgins, J., Perreault, L., Eckel, R. H., & Wright, K. P. (2013). Impact of insufficient sleep on total daily energy expenditure, food intake, and weight gain. Proceedings of the National Academy of Sciences, 110(14), 5695–5700. 10.1073/pnas.1216951110

45. Mathew, G. M., Reichenberger, D. A., Master, L., Buxton, O. M., Hale, L., & Chang, A.-M. (2022). Worse sleep health predicts less frequent breakfast consumption among adolescents in a micro-longitudinal analysis. The International Journal of Behavioral Nutrition and Physical Activity, 19(1), 70. 10.1186/s12966-022-01265-5

46. McNeil, J., Forest, G., Hintze, L. J., Brunet, J.-F., Finlayson, G., Blundell, J. E., & Doucet, É. (2017). The effects of partial sleep restriction and altered sleep timing on appetite and food reward. Appetite, 109, 48–56. 10.1016/j.appet.2016.11.020

47. Mela, D. J. (2006). Eating for pleasure or just wanting to eat? Reconsidering sensory hedonic responses as a driver of obesity. Appetite, 47(1), 10–17. 10.1016/j.appet.2006.02.006

48. Melanson, K. J., Westerterp-Plantenga, M. S., Campfield, L. A., & Saris, W. H. (1999). Blood glucose and meal patterns in time-blinded males, after aspartame, carbohydrate, and fat consumption, in relation to sweetness perception. The British Journal of Nutrition, 82(6), 437–446.

49. Monrroy, H., Borghi, G., Pribic, T., Galan, C., Nieto, A., Amigo, N., Accarino, A., Correig, X., & Azpiroz, F. (2019). Biological Response to Meal Ingestion: Gender Differences. Nutrients, 11(3), 702. 10.3390/nu11030702

50. Monteleone, P., Piscitelli, F., Scognamiglio, P., Monteleone, A. M., Canestrelli, B., Di Marzo, V., & Maj, M. (2012). Hedonic eating is associated with increased peripheral levels of ghrelin and the endocannabinoid 2-arachidonoyl-glycerol in healthy humans: A pilot study. The Journal of Clinical Endocrinology and Metabolism, 97(6), E917–924. 10.1210/jc.2011-3018

51. Najjar, R. P., Prayag, A. S., & Gronfier, C. (2024). Melatonin suppression by light involves different retinal photoreceptors in young and older adults. Journal of Pineal Research, 76(1). 10.1111/jpi.12930

52. Nakamura, K., & Nakamura, Y. (2018). Hunger and Satiety Signaling: Modeling Two Hypothalamomedullary Pathways for Energy Homeostasis. BioEssays: News and Reviews in Molecular, Cellular and Developmental Biology, 40(8), e1700252. 10.1002/bies.201700252

53. Paoli, A., Tinsley, G., Bianco, A., & Moro, T. (2019). The Influence of Meal Frequency and Timing on Health in Humans: The Role of Fasting. Nutrients, 11(4), 719. 10.3390/nu11040719

54. Pélissier, L., Lambert, C., Stensel, D. J., Beraud, D., Finlayson, G., Pereira, B., Boirie, Y., Duclos, M., Isacco, L., & Thivel, D. (2024). Individual variability and consistency of post-exercise energy and macronutrient intake, appetite sensations, and food reward in healthy adults. Appetite, 200, 107568. 10.1016/j.appet.2024.107568

55. Peschel, S. K. V., Tylka, T. L., Williams, D. P., Kaess, M., Thayer, J. F., & Koenig, J. (2018). Is intuitive eating related to resting state vagal activity? Autonomic Neuroscience: Basic & Clinical, 210, 72–75. 10.1016/j.autneu.2017.11.005

56. Polito, R., Valenzano, A., Monda, V., Cibelli, G., Monda, M., Messina, G., Villano, I., & Messina, A. (2022). Heart Rate Variability and Sympathetic Activity Is Modulated by Very Low-Calorie Ketogenic Diet. International Journal of Environmental Research and Public Health, 19(4), 2253. 10.3390/ijerph19042253

57. Prayag, A. S., Münch, M., Aeschbach, D., Chellappa, S. L., & Gronfier, C. (2019). Light Modulation of Human Clocks, Wake, and Sleep. Clocks & Sleep, 1(1), 193–208. 10.3390/clockssleep1010017

58. Qian, J., Morris, C. J., Caputo, R., Garaulet, M., & Scheer, F. A. (2019). Ghrelin is Impacted by the Endogenous Circadian System and by Circadian Misalignment in Humans. International Journal of Obesity (2005), 43(8), 1644–1649. 10.1038/s41366-018-0208-9

59. Rahman, S. A., Wright, K. P., Lockley, S. W., Czeisler, C. A., & Gronfier, C. (2019). Characterizing the temporal Dynamics of Melatonin and Cortisol Changes in Response to Nocturnal Light Exposure. Scientific Reports, 9, 19720. 10.1038/s41598-019-54806-7

60. Roenneberg, T., Wirz-Justice, A., & Merrow, M. (2003). Life between clocks: Daily temporal patterns of human chronotypes. Journal of Biological Rhythms, 18(1), 80–90. 10.1177/0748730402239679

61. Rogan, M. M., & Black, K. E. (2022). Dietary energy intake across the menstrual cycle: A narrative review. Nutrition Reviews, 81(7), 869–886. 10.1093/nutrit/nuac094

62. Ruddick-Collins, L. C., Byrne, N. M., & King, N. A. (2019). Assessing the influence of fasted and postprandial states on day-to-day variability of appetite and food preferences. Physiology & Behavior, 199, 219–228. 10.1016/j.physbeh.2018.11.015

63. Sargent, C., Zhou, X., Matthews, R. W., Darwent, D., & Roach, G. D. (2016). Daily Rhythms of Hunger and Satiety in Healthy Men during One Week of Sleep Restriction and Circadian Misalignment. International Journal of Environmental Research and Public Health, 13(2). 10.3390/ijerph13020170

64. Scheer, F. A. J. L., Hilton, M. F., Mantzoros, C. S., & Shea, S. A. (2009). Adverse metabolic and cardiovascular consequences of circadian misalignment. Proceedings of the National Academy of Sciences of the United States of America, 106(11), 4453–4458. 10.1073/pnas.0808180106

65. Scheer, F. A. J. L., Morris, C. J., & Shea, S. A. (2013). The Internal Circadian Clock Increases Hunger and Appetite in the Evening Independent of Food Intake and Other Behaviors. Obesity (Silver Spring, Md.), 21(3), 421–423. 10.1002/oby.20351

66. Schwartz, M. W., Woods, S. C., Porte, D., Seeley, R. J., & Baskin, D. G. (2000). Central nervous system control of food intake. Nature, 404(6778), 661–671. 10.1038/35007534

67. Solomon, S. J., Kurzer, M. S., & Calloway, D. H. (1982). Menstrual cycle and basal metabolic rate in women. The American Journal of Clinical Nutrition, 36(4), 611–616. 10.1093/ajcn/36.4.611

68. Spiegel, K., Leproult, R., & Van Cauter, E. (1999). Impact of sleep debt on metabolic and endocrine function. Lancet (London, England), 354(9188), 1435–1439. 10.1016/S0140-6736(99)01376-8

69. Stanić, Ž., Pribisalić, A., Bošković, M., Bućan Cvitanić, J., Boban, K., Bašković, G., Bartulić, A., Demo, S., Polašek, O., & Kolčić, I. (2021). Does Each Menstrual Cycle Elicit a Distinct Effect on Olfactory and Gustatory Perception? Nutrients, 13(8), 2509. 10.3390/nu13082509

70. Stevenson, R. J., Mahmut, M., & Rooney, K. (2015). Individual differences in the interoceptive states of hunger, fullness and thirst. Appetite, 95, 44–57. 10.1016/j.appet.2015.06.008

71. Tucker, J. A. L., McCarthy, S. F., Bornath, D. P. D., Khoja, J. S., & Hazell, T. J. (2025). The Effect of the Menstrual Cycle on Energy Intake: A Systematic Review and Meta-analysis. Nutrition Reviews, 83(3), e866–e876. 10.1093/nutrit/nuae093

72. Van Cauter, E., Polonsky, K. S., & Scheen, A. J. (1997). Roles of Circadian Rhythmicity and Sleep in Human Glucose Regulation*. Endocrine Reviews, 18(5), 716–738. 10.1210/edrv.18.5.0317

73. Watanabe, K., Umezu, K., & Kurahashi, T. (2002). Human olfactory contrast changes during the menstrual cycle. The Japanese Journal of Physiology, 52(4), 353–359. 10.2170/jjphysiol.52.353

74. Woods, S. C., Seeley, R. J., Porte, D., & Schwartz, M. W. (1998). Signals that regulate food intake and energy homeostasis. Science (New York, N.Y.), 280(5368), 1378–1383. 10.1126/science.280.5368.1378

75. Yukie, M., Aoi, I., Mizuki, K., & Toshiyuki, Y. (2020). Change in appetite and food craving during menstrual cycle in young students. International Journal of Nutrition and Metabolism, 12(2), 25–30. 10.5897/IJNAM2019.0264

76. Zeitzer, J. M., Dijk, D. J., Kronauer, R., Brown, E., & Czeisler, C. (2000). Sensitivity of the human circadian pacemaker to nocturnal light: Melatonin phase resetting and suppression. The Journal of Physiology, 526 Pt 3(Pt 3), 695–702. 10.1111/j.1469-7793.2000.00695.x

